# Innate immune sensing of cell traversal by *Plasmodium* sporozoites drives protective T cell responses

**DOI:** 10.64898/2026.03.20.713171

**Authors:** Kai G. Pohl, Xin Gao, Jason McGowan, Parnika Mukherjee, Shirley Le, Patricia Carreira, Chinh Ngo, Henry J. Sutton, Sophia Brumhard, Anna Hiller, Hannah Kelly, Larissa Liow, Adrian Lo, Larissa Henze, Lucie Loyal, Rogerio Amino, Si Ming Man, Lynette Beattie, Leif Erik Sander, Ian A Cockburn

## Abstract

Attenuated *Plasmodium* sporozoites elicit robust adaptive immune responses that protect against parasite challenge, yet how they engage the innate immune system remains poorly understood. To obtain sterile preparation of sporozoites we used flow cytometry to isolate pure GFP^+^ *P. berghei* and *P. falciparum* sporozoites, which were sorted onto murine bone marrow–derived macrophages (BMDMs) or human PBMCs to assess innate immune activation. Sporozoites induced a distinctive activation pattern resembling cell wounding responses, consistent with their ability to traverse host cells en route to the liver. Traversal-deficient sporozoites failed to activate BMDMs and showed impaired priming of protective CD8^+^ T cell immunity. This wounding response was independent of inflammasome or Toll-like receptor signaling but required γδ T cells, which are known to support CD8^+^ T cell responses against *Plasmodium*. These findings reveal a previously unappreciated innate sensing mechanism triggered by cell traversal that underpins the potent immunogenicity of *Plasmodium* sporozoites.

## Main

Malaria remains a major global threat with nearly 250 million clinical disease episodes and an estimated 608000 deaths in 2022^1^. While symptomatic disease is caused by asexual blood stage parasites, most vaccines target the sporozoite (SPZ) stages of the parasite based on the observation that multiple injections with high doses of irradiated *Plasmodium* SPZ can induce sterile protection against subsequent live parasite challenge^2,3^. The current RTS,S/AS01 and R21/MatrixM vaccines aim to mimic a component of SPZ induced immunity by inducing antibodies targeting the circumsporozoite protein, sterile irradiated SPZ themselves are also being tested as a potential vaccine. In addition to antibodies SPZ are also capable of priming significant CD8^+^ T cell responses and ablation of CD8^+^ T-cells before challenge abrogates protection^4,5^. However, the strict requirement for SPZ to be viable during immunization and the fact that SPZ only develop inside of mosquitoes preclude scalable manufacturing and logistics of a potential SPZ vaccine^6^.

The basis for the potent immunogenicity of *Plasmodium* SPZ is not fully understood. The magnitude and quality of an adaptive immune response is largely determined by sensing of pathogen associated molecular patterns (PAMPs) through germline-encoded pattern recognition receptors (PRRs) of the innate immune system^7^. Upon pathogen sensing, multiple innate immune pathways and interactions of innate and adaptive immune cells encode the features of the induced adaptive response^8^. Many innate sensors and pathways are able to detect *Plasmodium* blood stage parasites including hemozoin^9^, which is unique to blood stages, and parasite nucleic acid^10^ and GPI anchors^11^ which are shared with SPZ. However, given that potent immune responses can be induced to the relatively small numbers of non-replicating parasites injected in the SPZ vaccine, it seems likely that SPZ have additional PAMPs not present in blood stages. Recent studies showed that CD8+ T cell responses to live attenuated SPZs are supported by innate-like gamma delta T cells though the signals driving these responses are unknown^12,13^.

To better understand the innate immune recognition of *Plasmodium* SPZ, we developed a reductionist system in which mouse or human innate immune cells were incubated with sterile, flow-sorted SPZ or blood stage parasites. Bulk and single cell RNA sequencing revealed that SPZ triggered a distinct transcriptional activation that was characterized by a conserved plasma membrane damage response. During their migration from the skin to the liver, SPZ traverse and wound multiple cell types before establishing infection in hepatocytes. Importantly, SPZ unable to traverse failed to induce protective CD8 T cell responses. We thus propose that this traversal-associated wounding—absent in blood-stage parasites—serves as a key driver of the potent immunogenicity of *Plasmodium* SPZ.

## Results

### FACS-Based Purification Enables Dissection of Macrophage Responses to Plasmodium s and Blood Stages

Attempts to dissect the innate response to *Plasmodium* SPZ are hampered by difficulties in differentiating parasite driven responses from those induced by mosquito material that contaminates sporozoite preparations. To overcome this, we used fluorescence-activated cell sorting (FACS) to separate *P. berghei* ConF^14^ SPZ expressing GFP under the control of the HSP70 promotor (referred to as control *Pb*SPZ) from mosquito material directly following sporozoite preparation from salivary glands (Fig. 1a). GFP expressing *Pb*SPZ were easily detectable by flow cytometry (Fig. 1b) and enumeration of *Pb*SPZ content within salivary gland material between different mosquito batches showed an expectedly high variation (Fig. 1c), thus underscoring the utility of sorting *Pb*SPZ from these preparations to standardize downstream experiments.

**Fig. 1.**
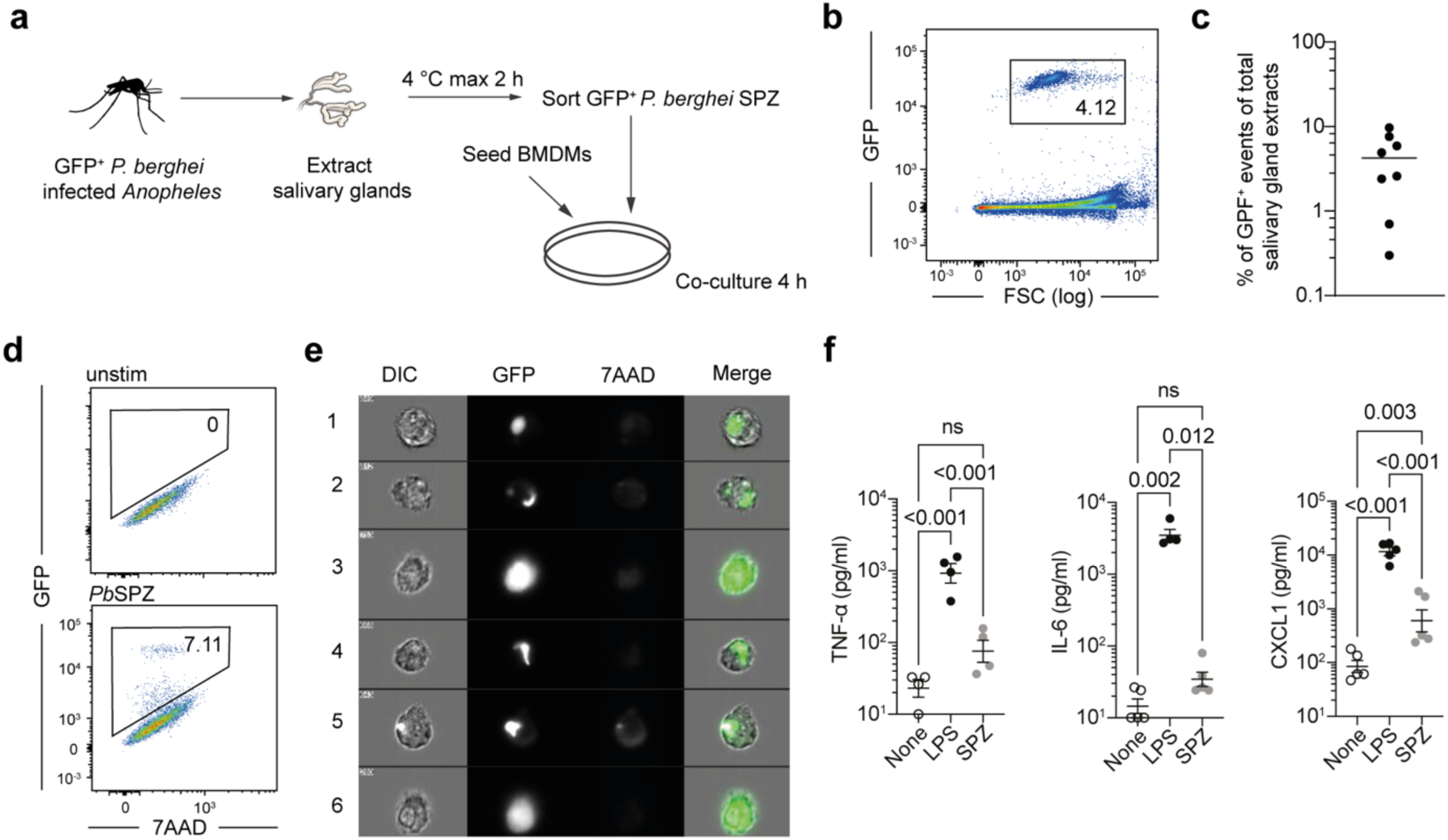
A reductionist system to probe innate immune responses to Plasmodium parasites. a, Schematic of experimental setup. **b,** GFP^+^ ConF Sporozoites (*Pb*SPZ) are clearly detectable in whole salivary gland extracts. **c,** Frequency of *Pb*SPZ within salivary gland extracts vary across orders of magnitude. **d,** A population of GFP^+^ macrophages is present after 4 h of co-culture with *Pb*SPZ. **e,** Imaging cytometry of GFP^+^ macrophages after 4 h of co-culture with *Pb*SPZ identifies 2 subtypes of interactions: panels 3 & 6: diffuse GFP; panels 1,2,4,5: intact SPZ inside or attached to the surface of macrophages. **f,** TNF-α, IL-6 and CXCL1 levels in the supernatant after 24 hours of co-culture with indicated *Pb*SPZ at an MoI of 0.5; data shown are experimental replicates from 2 independent experiments for each condition, mean ± SD shown, analysis via linear mixed model.

To assess innate immune responses to purified *Pb*SPZ in a reductionist system we sorted 5x10^4^ *Pb*SPZ directly onto 1x10^5^ primary bone marrow derived macrophages (BMDMs) at multiplicity of infection (MoI) of 0.5 (Fig. 1a). Time lapse imaging revealed that shortly after sorting, SPZ were viable and displayed motile gliding over the surface of the macrophage layer, with some evidence of uptake by macrophages even at this early timepoint (Supplementary video 1). After 4 hours of co-culture, GFP^+^ macrophages were evident by flow cytometry (Fig. 1d) and image flow analysis of GFP^+^ events confirmed these as macrophages with associated SPZ (Fig. 1e). Two distinct types of GFP^+^ macrophages were evident, those with diffuse GFP signal (Fig. 1e, 3 & 6) and those with seemingly intact SPZ present inside or attached to a macrophage (Fig. 1e 1,2,4 & 5). However, despite clear evidence of interactions between *Pb*SPZ and BMDMs we were unable to detect significant production of key inflammatory cytokines IL-6 and TNF-α (Fig. 1f). Production of CXCL1 was around 30-fold lower than that induced by LPS stimulation (Fig. 1f). We also determined that blood stage parasites (*Pb*RBCs) could be purified in an analogous system to allow for direct side-by-side comparison of innate responses to the two life cycle stages (Extended data Fig. 1a). Similarly to *Pb*SPZ, GFP^+^ *Pb*RBCs could be readily detected in the blood of infected mice and separated from uninfected cells by FACS (Extended data Fig. 1b). In the same co-culture system (Extended data Fig. 1c), these GFP^+^ *Pb*RBCs were taken up by BMDMs (Extended data Fig. 1d) and were readily detectable by image flow (Extended data Fig. 1e).

**Fig. 2:**
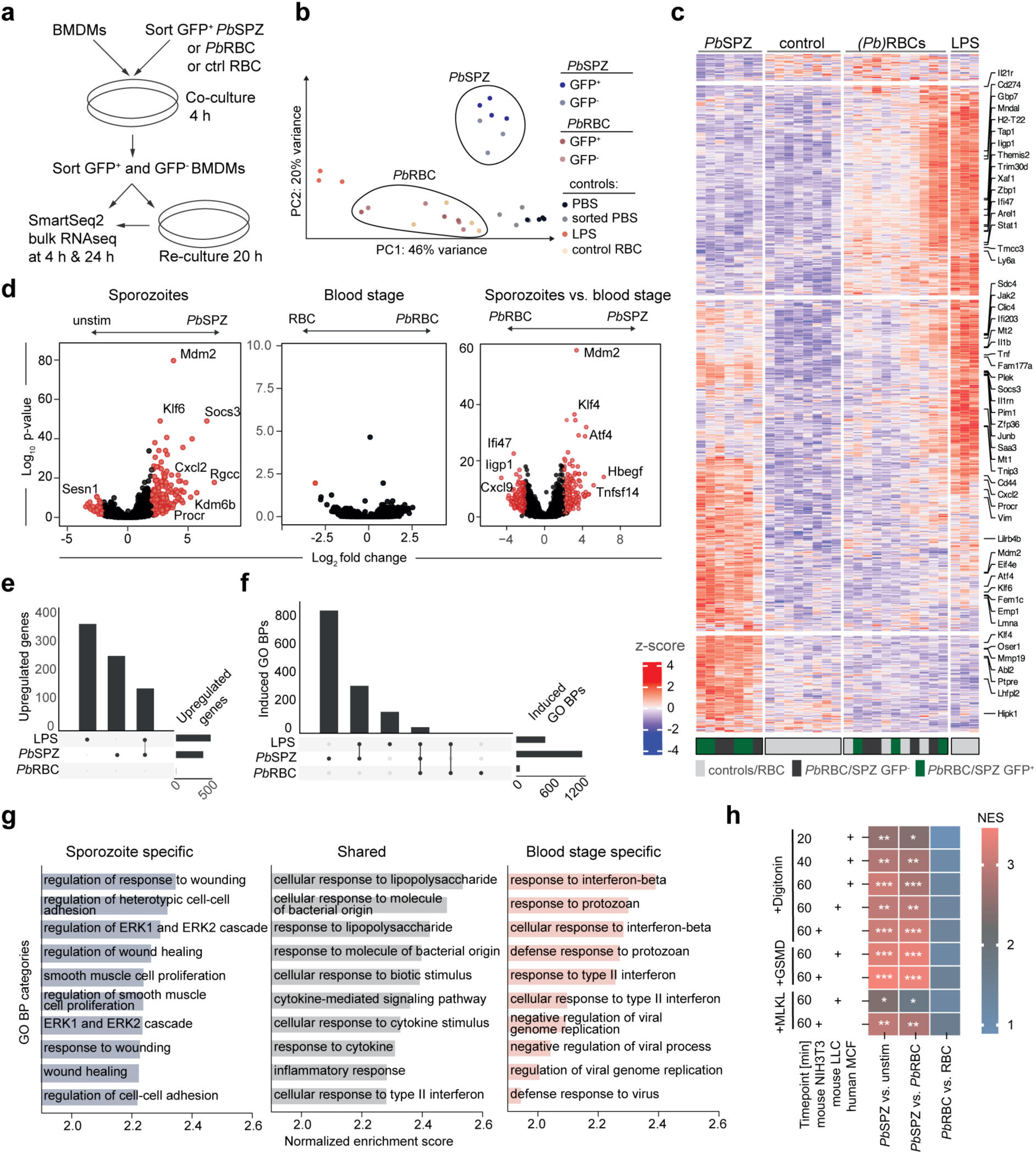
Transcriptional profiling of BMDMs reveals distinct activation patterns induced by sporozoite and blood stage Plasmodium parasites. a, Schematic of experimental setup. **b,** Principal Component Analysis (PCA) clearly separates BMDMs co-cultured with *Pb*SPZ and *Pb*RBC. **c,** Volcano plots of differentially expressed genes between the indicated conditions irrespective of GFP-status showing unique transcriptional signatures induced by *Pb*SPZ and *Pb*RBC. **d,** Upset plot of upregulated genes through co-culture with indicated stimuli showing quantitative and qualitative differences in transcriptional activation induced by LPS, *Pb*SPZ and *Pb*RBC. **e,** Upset plot showing unique and overlapping gene ontology (GO) biological processes (BP) enriched in transcriptional signatures induced by indicated stimuli. **f,** Heatmap of differentially expressed genes with key genes indicated. **g,** GO BPs specifically induced by *Pb*SPZ, *Pb*RBC or shared shows a wounding response and ERK1 and ERK2 cascade signature specific for *Pb*SPZ co-cultured BMDMs. **h,** Gene set enrichment analysis (GSEA) showing highly significant enrichment of curated plasma membrane injury signatures^21,22^ specifically in *Pb*SPZ treated BMDMs. Enrichment was calculated based on genes ordered by log2 fold change between the on top indicated conditions. Transcriptomic signatures were generated by subjecting indicated cell lines to sublethal doses of indicated stimuli and assessing DEGs to untreated controls at indicated timepoints. *** = p < 1x10^-25^, ** = p < 1x10^-10^, * = p < 5x10^-2^.

**Fig. 3:**
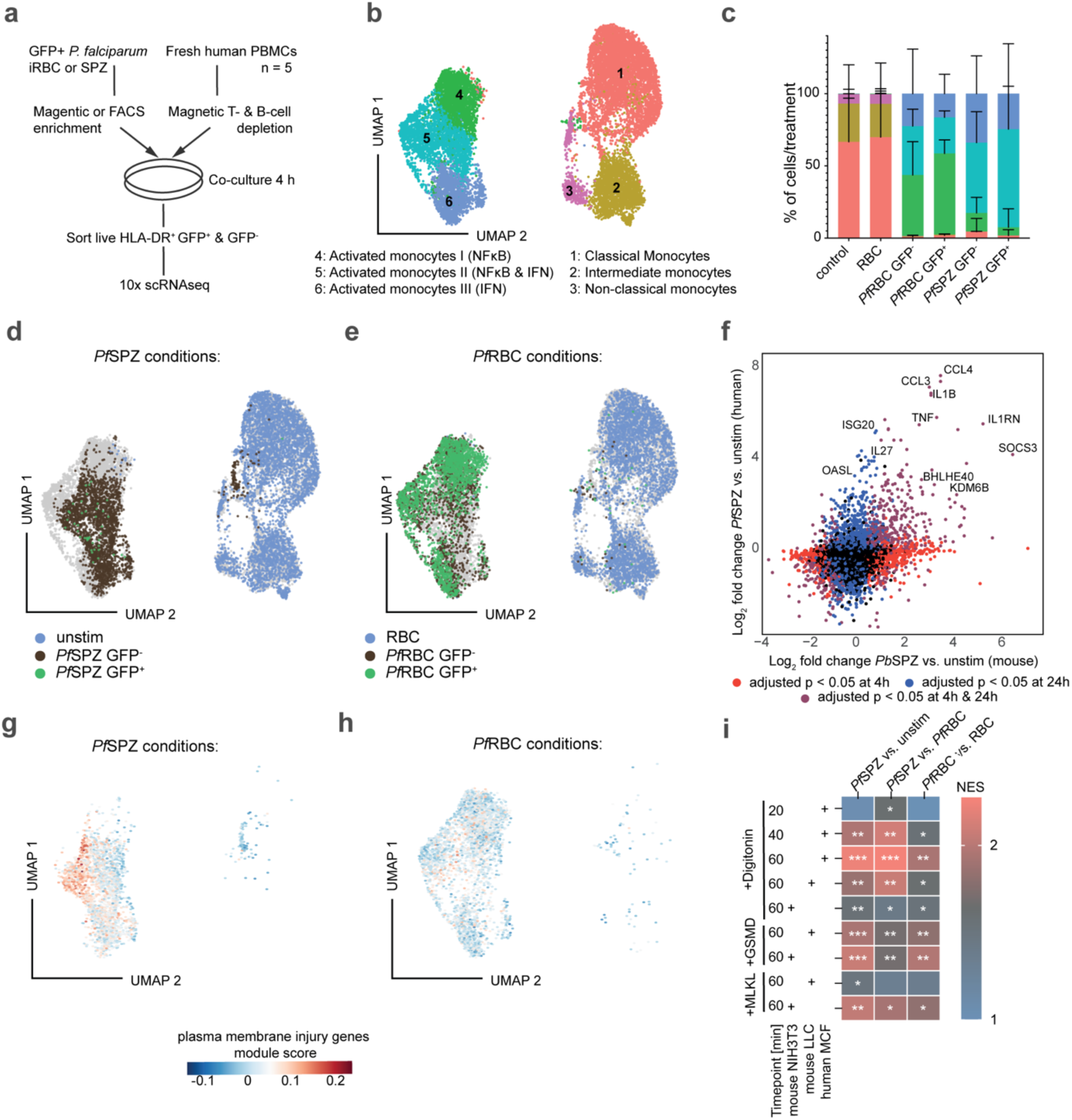
Sporozoite-induced activation of human monocytes shows cross-species overlap and a plasma membrane injury signature. a, Schematic of experimental setup. **b,** UMAP plot of primary human monocytes stimulated with *Pf*SPZ, *Pf*RBC or control RBC. **c,** Cell type distribution across conditions (mean + sd 5 donors across two independent experiments shown). **d,** Cells from *Pf*SPZ stimulated and control conditions highlighted on UMAP. **e,** Cells from *Pf*RBC and control RBC stimulated conditions highlighted on UMAP. **f,** Log2 fold change of genes of *Pf*SPZ stimulated monocytes vs. unstimulated control (y-axis) and *Pb*SPZ stimulated BMDMs vs. unstimulated control (x-axis) showing a conserved SPZ specific innate response. Genes were mapped between species based on ensembl annotation. **g,h,** UMAP depicting module score for combined plasma membrane injury gene sets for cells co-cultured with *Pf*SPZ (G) and *Pf*RBC (H). **i,** GSEA (as in Fig. 2) showing strongest enrichment of plasma membrane injury genes in *Pf*SPZ vs. *Pf*RBC contrast. *** = p < 1x10^-10^, ** = p < 1x10^-5^, * = p < 5x10^-2^.

**Fig. 4:**
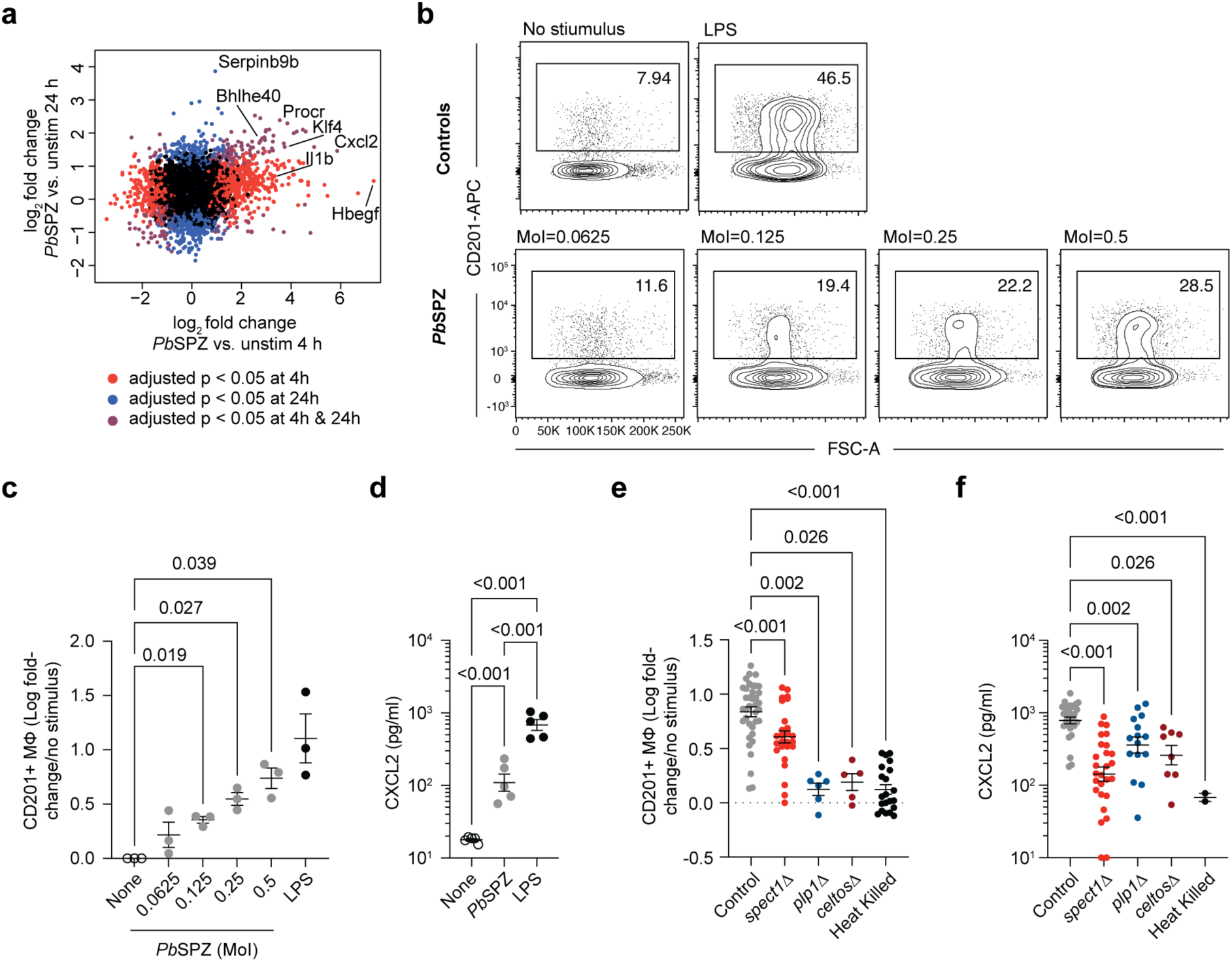
SPZ induce activation of BMDMs in a CT dependent manner. a, Scatter plot depicting gene expression log2 fold changes in BMDMs stimulated with *Pb*SPZ for 4 h (x-axis) and 24 h (y-axis) compared to control, colored by significance levels as indicated, showing that Procr (coding for CD201) and Cxcl2 are consistently upregulated over both timepoints. **b,** Representative flow cytometry plots showing upregulation of CD201 on the surface of BMDMs after 24 hours of incubation with LPS or *Pb*SPZ at the indicated MoIs. **c,** Quantification of data from A; data from a single experiment, mean ± SD indicated, analysis via one-way ANOVA with Dunnett’s multiple comparisons test. **d,** CXCL2 levels in the supernatant after 24 hours of co-culture with *Pb*SPZ at an MoI of 0.5; data shown are experimental replicates from 2 independent experiments, mean ± SD shown, analysis via linear mixed model. **e,** CD201 expression on BMDMs after 24 hours co-culture with the indicated sporozoites at an MoI of 0.5; data shown are technical replicates from at least 3 independent experiments per condition, mean ± SD shown, analysis via linear mixed model. **f,** CXCL2 levels in the supernatant after 24 hours of co-culture with indicated *Pb*SPZ at an MoI of 0.5; data shown are technical replicates from at least 2 independent experiments per condition, mean ± SD shown, analysis via linear mixed model.

**Fig. 5:**
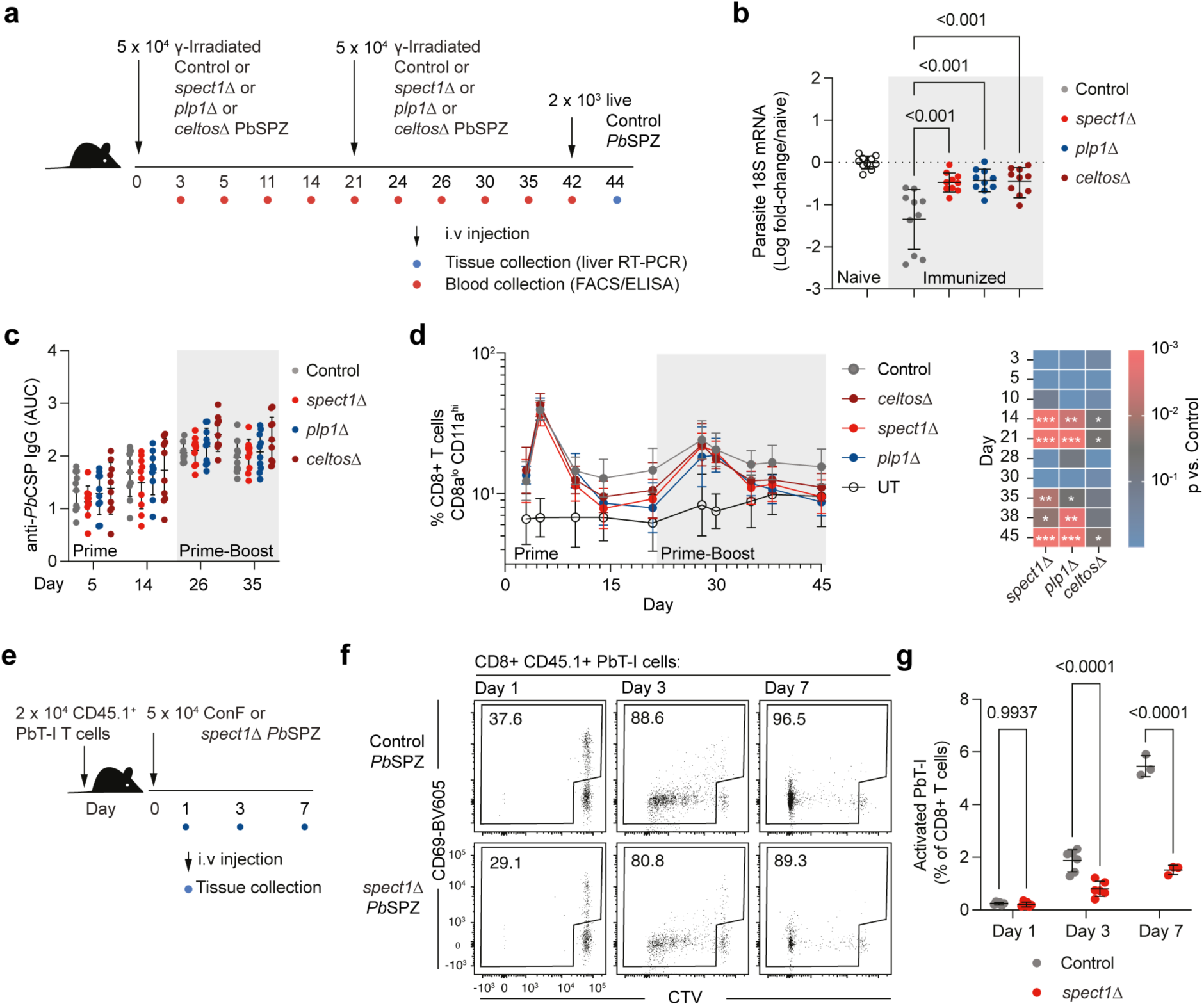
CT deficient irradiated sporozoites fail to induce optimal CD8+ T cell responses. a, Schematic of experiment to determine the induction of protective immunity by different CT deficient *Pb*SPZ. **b,** Parasite burden assessed by 18S RT-PCR in mice immunized and challenged as in A; data pooled from 2 independent experiments, mean ± SD shown, analysis via linear mixed model. **c,** Anti-*Pb*CSP IgG in the sera of mice immunized as in A; data pooled from 2 independent experiments, mean ± SD shown, analysis via linear mixed model, no significant differences were observed across any groups. **d,** Peripheral blood CD8^+^ T cell responses in mice immunized as in A; data pooled from 2 independent experiments, mean ± SD shown, analysis via linear mixed model, with p values for comparisons to the control group presented in the heatmap (* = p<0.05; ** = p<0.01, *** = p<0.001). **e,** Schematic of experiment to determine the activation of *Plasmodium*-specific PbT-I CD8^+^ T cells by CT deficient *Pb*SPZ. **f,** Representative flow cytometry plots showing the activation of PbT-I cells by Control and *spect1*Δ *Pb*SPZ. **g,** Quantification of data from E-F; data from 3 independent experiment, mean ± SD indicated, analysis via linear mixed model.

**Fig. 6:**
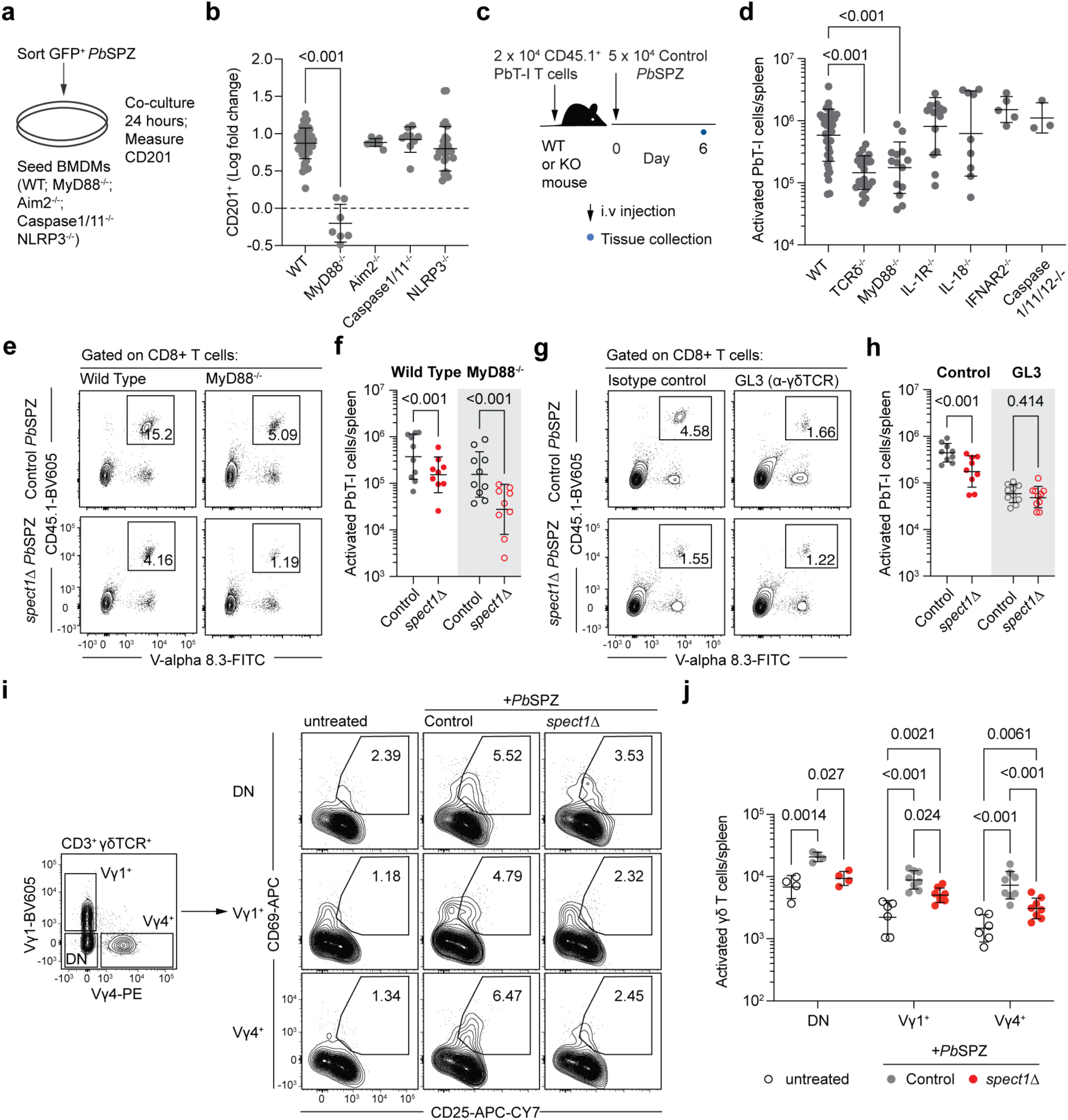
Sensing of CT requires gamma delta T cells. a, Schematic of approach to determine role of different innate immune pathways in the sensing of *Pb*SPZ. **b,** CD201 expression on BMDMs after 24 hours co-culture with the indicated sporozoites at an MoI of 0.5; data shown are techincal replicates from at least 2 independent experiments, mean ± SD shown, analysis via linear mixed model. **c,** Schematic of experiment to determine the activation of *Plasmodium*-specific PbT-I CD8^+^ T cells in mice lacking components of innate immune sensing pathways. **d,** Number of PbT-I cells in the spleen of mice immunized as in C data from at least 2 independent experiments per condition, mean ± SD indicated, analysis via linear mixed model. **e,** Representative flow cytometry plots showing the activation of PbT-I cells in wild type and MyD88^-/-^ mice by Control and *spect1*Δ *Pb*SPZ **f,** Quantification of data from E; data from 2 independent experiments, mean ± SD indicated, analysis via linear mixed model. **g,** Representative flow cytometry plots showing the activation of PbT-I cell by control and *spect1*Δ *Pb*SPZ in control mice or mice receiving GL3 mAb (anti-γδTCR). **h,** Quantification of data from E; data from 2 independent experiments, mean ± SD indicated, analysis via linear mixed model. **I,** Activation of γδT cell subsets in spleens of mice 24 h after immunization with wt or *spect1*Δ *Pb*SPZ or control mice. **j,** Quantification of J. Results of 2 independent experiments shown for Vγ1 and Vγ4 T cells and 1 experiment of double negative (DN) γδT cells, statistical analysis via 2-way ANOVA with Tukey correction.

### *Pb*SPZ induce a unique pattern of macrophage activation

To unbiasedly assess innate activation by the *Plasmodium* life cycle stages, we profiled BMDM transcriptional responses after 4 h of incubation with *Pb*SPZ or *Pb*RBC at an MoI of 0.5 or with LPS (Fig. 2a). Because low numbers of *Pb*SPZ were limiting the scale of our experiments, we adapted the SmartSeq2 single cell RNA-seq protocol and sorted 50 cells per replicate and condition with slightly adjusted lysis buffer conditions (see Methods). This bulk RNA-seq approach allowed us to multiplex conditions and to separate GFP^+^ from GFP^-^BMDMs before RNA extraction (Fig. 2a). To control for potential contaminants from parasite sorting, we included a ‘sorted PBS’ control in which BMDMs were treated with equal volumes of PBS that had passed through the sorter. In addition, to profile long-lasting parasite-induced transcriptional changes in BMDMs, we re-cultured sorted GFP^+^ and GFP^-^BMDMs in absence of additional stimuli for 20 h before RNA sequencing (Fig. 2a).

Principal components analysis (PCA) of RNA-seq data clearly separated BMDM transcriptional responses between blood stage and SPZ conditions at 4 h (Fig. 2b), but not at 24 h (Extended data Fig. 2b), revealing strikingly divergent responses to the different life cycle stages at the early timepoint that converged onto a shared transcriptional program over time, while LPS induced transcriptional changes appeared distinct from parasite induced responses at both time points. Interestingly, after 4 h of co-culture, transcriptional responses in GFP^+^ and GFP^-^ BMDMs were highly similar within BMDMs co-cultured with either life cycle stage (Fig. 2b and Extended data Fig. 2c), indicating that we likely picked up secondary transcriptional responses to cytokines produced early during co-culture. Alternatively, this may suggest that transcriptional changes were principally the result of exposure to parasites rather than parasite phagocytosis.

We next looked at differentially expressed genes (DEGs) between key conditions irrespective of GFP status at 4 h (Supplementary data 1). Unsupervised clustering of these genes revealed 3 main clusters of genes (Fig. 2c): A cluster of genes that were jointly upregulated upon either treatment – though to varying degrees –, a cluster that was unique to *Pb*SPZ treated BMDMs and a last cluster of genes that were induced upon LPS and *Pb*RBC – but not by *Pb*SPZ stimulation (Fig. 2d). Direct comparison of expression levels between BMDMs co-cultured with *Pb*RBC and *Pb*SPZ revealed an abundance of genes with life cycle stage-specific regulation (Fig. 2d). Quantitatively, *Pb*SPZ and LPS induced similar numbers of upregulated genes (Fig. 2e) while, surprisingly, *Pb*RBC co-culture did not induce significant upregulation of any gene when contrasted against the RBC control condition (Fig. 2d & e). We thus show quantitative and qualitative differences in innate cell activation capacity of the two *Plasmodium* life cycle stages.

We next used gene set enrichment analysis (GSEA)^15,16^ to assess transcription factor binding sites around regulated genes and parasite-induced biological processes (BPs) as annotated by gene ontology (GO)^17,18^. This analysis revealed that both life cycle stages induced NFκB-driven gene expression in BMDMs (Extended data Fig. 2d). However, *Pb*RBCs uniquely induced IRF driven expression (Extended data Fig. 2d), while *Pb*SPZ uniquely induced *Bach1, Bach2, AP1* and *Stat5* driven expression (Extended data Fig. 2d); which have been implicated as key players in macrophage polarization^19^ and oxidative stress responses^20^. *Pb*SPZ induced the largest number of GO BPs (1200) of which 300 were shared between LPS- and 50 between LPS and *Pb*RBC stimulated BMDMs, respectively (Fig. 2f, Extended data Fig. 2e). Looking at shared and unique GO BPs induced by co-culture with *Pb*SPZ and *Pb*RBCs revealed that *Pb*SPZ stimulation specifically induced a ‘wounding response’ signature as well as ERK1 and ERK2 pathways (Fig. 2g). Plasma membrane disruption has previously been implicated as a driver of inflammatory responses^21,22^. As such, we hypothesised that sensing of plasma membrane damage could partly explain the BMDM activation phenotype. To test this, we manually curated gene sets of plasma membrane disruption response genes from experiments wherein cells were subjected to sub-lethal plasma membrane damage and transcriptionally profiled after a short recovery phase (Supplementary data 1)^21,22^. Strikingly, all curated gene sets were highly enriched in BMDMs co-cultured with *Pb*SPZ when compared to control or *Pb*RBC co-cultured BMDMs but not enriched in BMDMs co-cultured with *Pb*RBC (Fig. 2h). We thus identify membrane disruption as a likely driver of *Pb*SPZ specific transcriptional responses in innate immune cells.

### Plasmodium falciparum SPZ induce a wounding response in human monocytes

We next wanted to know if our findings with *Pb*SPZ and BMDMs were generalizable to primary human innate immune cells interacting with human infecting *Plasmodium falciparum* SPZ (*Pf*SPZ). Accordingly, NF54-CGL *Pf*SPZ^23^, expressing a GFP-luciferase fusion protein under the control of the CSP promotor, were isolated from *An stephensi* salivary glands, sort-purified by flow cytometry (Extended data Fig. 3a) and co-cultured with primary human T and B cell depleted PBMCs at an MOI of 0.5 (Fig. 3a). Additionally, we co-cultured T and B cell depleted PBMCs from the same donors with magnetically enriched *P. falciparum* infected erythrocytes expressing GFP (*Pf*RBC) and control uninfected erythrocytes (RBC). Because of the marginal stimulation observed from *Pb*RBC, we co-cultured *Pf*RBC with human innate cells at an MoI of 5. As with murine BMDMs, we were readily able to identify GFP^+^ cells within the HLA-DR^+^ cell compartment using flow cytometry after 4 hours of co-culture (Extended data Fig. 3b). Analogously to our previous experiments, we separated GFP^+^ from GFP^-^ cells using FACS after 4 hours of co-culture and subjected these populations to single cell RNA sequencing (Fig. 3a).

Uniform Manifold Approximation and Projection (UMAP) clustering of scRNA-seq transcriptomic data after removing small numbers of contaminating T and B cells as well as a small cluster of dendritic cells, identified two main clusters of monocytes, of which one consisted predominantly of resting cells from the mock or uninfected RBC conditions, while the other exclusively consisted of activated cells from the *Pf*SPZ and *Pf*RBC conditions (Fig. 3b). Manual annotation of the three monocyte subclusters for the unstimulated and RBC conditions revealed classical, intermediate and non-classical monocytes based on expression of canonical monocyte marker genes (Extended data Fig. 3c). Activated monocytes from the *PfRBC* and *Pf*SPZ conditions also fell into 3 subclusters which were annotated as activated monocytes (AM) I, II and III (Extended data Fig. 3c, Supplementary data 2). Marker gene signatures indicated mostly NF-κB-driven gene expression in cluster AM I, cluster AM 3 was mainly characterized by type-I IFN response genes, while cluster AM 2 contained a combined NF-κB and type I IFN signatures. Interestingly, cells stimulated with *Pf*SPZ fell more frequently into monocyte cluster 5 (Fig. 3c and d), while cells stimulated with *Pf*RBC fell into cluster 4 (Fig. 3 c and e).

We next asked how our two model systems compared on the transcriptional level. Using pseudo bulk^24^ analysis of the human single cell dataset, we performed the same pairwise analyses between treatment conditions as previously in the mouse/*P.berghei* model. Human monocytes regulated large numbers of pro-inflammatory mediators after co-culture with either parasite stage, like cytokines IL1B, TNF and chemokines CCL2 and CCL3 (Extended data Fig. 4a). Strikingly, sporozoite stimulation induced the upregulation of 99 shared genes between the species (Fig. 3f, OR: 3,7, p < 0.01 Fisher’s exact test). Using slightly less stringent cut-offs (log2 fold change: <1,5 and adjusted p: <0.05), we identified a shared module of 260 genes that were upregulated in *Pb*SPZ*, Pf*SPZ and *Pf*RBC as well as 70 genes that were upregulated in *Pb*SPZ and *Pf*SPZ stimulated innate cells, respectively (Extended data Fig. 4b). Overrepresentation analysis (ORA) of this shared expression module revealed key inflammatory pathways (TNF IFN-y, IL-6 signaling) but also Hypoxia and Apoptosis as enriched terms (Extended data Fig. 4c).

We next asked if the plasma membrane injury signature observed in the *P. berghei* – BMDM experiments was also enriched in *P. falciparum* stimulated human monocytes. Strikingly, cells stimulated with *Pf*SPZ showed stronger enrichment of the plasma membrane injury genes (Fig. 3g) than cells stimulated with *Pf*RBC (Fig. 3h). In analogy to our earlier analysis, we also tested gene set enrichment of each individual plasma membrane injury gene set. While *Pf*SPZ and *Pf*RBC stimulated monocytes both showed enrichment of these gene sets when compared to their respective controls, highest enrichment was apparent when genes were ranked by expression between *Pf*SPZ and *Pf*RBC condition (Fig. 3i), thus confirming that *Pf*SPZ specifically induce a plasma membrane damage response.

### CT deficient parasites fail to induce BMDM activation

To mechanistically interrogate *Pb*SPZ sensing of BMDMs through measurement of cell surface or secreted molecules, we analysed our bulk RNAseq dataset for genes that were consistently upregulated at 4 h and 24 h timepoints. Interestingly, *Cxcl2*, a key inflammatory marker of tissue injury^25^ was consistently upregulated (Fig. 4a). In addition, *Procr* (encoding protein C receptor or CD201) was also upregulated across timepoints. CD201 has been implicated in presenting ligands to γδ T cells^26,27^ which have a role in promoting CD8^+^ T-cell responses to sporozoite vaccination^12^. As expected, we detected dose-dependent expression of CD201 by flow cytometry (Fig. 4b,c; Extended data Fig. 5a) and confirmed presence of CXCL2 in the supernatants of BMDMs co-cultured with *Pb*SPZ via ELISA (Fig. 4d). Interestingly, despite high expression of *IL1B/Il1b* in human monocyte-*Pf*SPZ and mouse BMDM-*Pb*SPZ experiments (Fig. 3f), IL-1β was not secreted unless BMDMs were pre-primed with LPS (Extended data Fig. 5b). IL-1β secretion follows inflammasome activation which requires a priming and an activation signal. Thus, our data indicates that *Pb*SPZ can constitute signal 2 necessary for inflammasome activation but are themselves not able to prime inflammasome activation (signal 1)^28^.

Our RNA-seq data showed that *Pb*SPZ, but not blood stages, induced a cell wounding transcriptional response in macrophages. Prior to productive invasion of hepatocytes, SPZ have been shown to undertake cell traversal (CT) of macrophages, endothelial cells and hepatocytes en route to the final target of infection^29^. CT is mediated by a family of proteins including Sporozoite Microneme Protein Essential for Cell Traversal (SPECT1)^30^, Perforin like protein 1 (PLP1; also known as SPECT2)^31^ and Cell-traversal Protein for Ookinetes and Sporozoites (CelTOS)^32^. Notably PLP1 carries a membrane attack complex domain making it a strong candidate as a mediator of cell wounding^31^, while CelTOS, which is a *Plasmodium* specific gene also forms pores has the ability to damage cell membranes^33^. Accordingly, GFP^+^ SPECT1 KO (*spect1*Δ), GFP^+^ PLP1 KO (*plp1*Δ) and GFP^+^ CelTOS KO (*celtos*Δ) and control fluorescent *Pb*SPZ ^14,32,34^ were sorted onto BMDMs followed by measurement of CXCL2 secretion and CD201 upregulation after 24 hours. As an additional control, we examined the activation of BMDMs by heat-killed control parasites. All CT–deficient mutant parasites induced lower CXCL2 secretion and reduced CD201 upregulation compared to traversal-competent control parasites (Fig. 4e,f). Notably, *spect1Δ* parasites also caused less cell wounding in BMDMs, as indicated by reduced Rhodamine-Dextran uptake (Extended data Fig. 5c,d). Importantly, since all three CT-deficient parasites showed similar phenotypes in BMDMs and heat-killed parasites failed to activate BMDMs, it is likely that the process of CT itself—rather than the specific presence of SPECT1, PLP1, or CelTOS— is sensed by BMDMs.

### CT deficient PbSPZ fail to elicit protective CD8^+^ T cell responses

Given the reduced macrophage activation by CT-deficient *Pb*SPZ, we next asked whether these parasites also induced impaired adaptive immune responses. Mice were immunized twice at three-week intervals with either *spect1*Δ, *plp1*Δ, *celtos*Δ, or control *Pb*SPZ. Blood was collected at defined time points following each immunization for the analysis of antibody and CD8^+^ T cell responses (Fig. 5a; Extended data Fig. 6a). Three weeks after the second immunization, mice were challenged with control *Pb*SPZ, and parasitemia was assessed by RT-PCR (Fig. 5a). While all immunized mice were protected against infection, the degree of protection was significantly reduced in mice immunized with traversal-deficient *Pb*SPZ compared to controls (Fig. 5b).

Protection against sporozoite challenge is generally attributed to antibodies targeting surface antigens, notably the circumsporozoite protein^35^, and to CD8^+^ T cells capable of eliminating infected hepatocytes^36–38^. Antibody responses to *P. berghei* circusmsporozoite protein (*Pb*CSP) were comparable between mice immunized with traversal-deficient and control *Pb*SPZ for IgG (Fig. 5c) and IgM (Extended data Fig. 6b). CD8^+^ T cell responses were also initially similar when assessed in the blood by CD11a upregulation and CD8a downregulation^39^. However, by day 14, CD8^+^ T cell responses in mice immunized with traversal-deficient *Pb*SPZ had declined to background levels, while responses remained detectable in mice immunized with control *Pb*SPZ (Fig. 5d).

Impaired CD8^+^ T cell activation following immunization with traversal-deficient parasites may result from reduced innate immune activation within lymphoid tissues, where T cell priming occurs^40,41^. Alternatively, reduced responses could reflect inefficient liver infection and impaired presentation of late liver-stage antigens^42^. To investigate the site and timing of priming, we analyzed the activation of CellTrace Violet-labelled *P. berghei* specific PbT-I TCR transgenic CD8^+^ T cells in the spleen at 24 hours, 3 days, and 7 days following irradiated *Pb*SPZ inoculation (Fig. 5e). As early as 24 hours post-immunization, PbT-I cells primed by *spect1*Δ *Pb*SPZ exhibited reduced expression of the early activation marker CD69 compared to those primed by control *Pb*SPZ (Fig. 5f). By 3 days post-immunization, the number of activated PbT-I cells was significantly reduced in mice immunized with *spect1*Δ *Pb*SPZ, with this impairment becoming more pronounced by day 7 (Fig. 5g).

Importantly, the observed defect in CD8^+^ T cell activation did not appear to result from altered splenic tropism. Although *spect1*Δ *Pb*SPZ exhibited impaired liver infection, as reported previously, their distribution to the spleen was comparable to that of wild-type parasites (Extended data Fig. 6c). This conclusion was further supported by the observation that antibody responses, which are also primed in the spleen^43^, remained intact following immunization with traversal-deficient parasites (Fig. 5c and Extended data Fig. 6b).

### Gamma-delta T cells sense CT to potentiate CD8^+^ T cell responses

Finally, we sought to identify the innate immune pathways responsible for sensing *Pb*SPZ by analysing CD201 expression of BMDMs deficient in a range of innate sensing and signaling molecules covering the inflammasome (*Nlrp3^-/-^, Aim2^-/-^, Caspase1/11^-/-^*) and TLR signalling pathways (*MyD88^-/-^*) (Fig. 6a). Notably, CD201 upregulation occurred normally in BMDMs from all knockout strains except those lacking MyD88, suggesting a requirement for this adaptor in sensing of *Pb*SPZ (Fig. 6b). To assess whether these innate pathways affect CD8^+^ T cell priming *in vivo*, we examined activation of PbT-I cells in various knockout mice (Fig. 6c). In addition to mice lacking TLR signaling (*Myd88*^-/-^) and inflammasome components (*Il1r1*^-/-^, *Il18r1*^-/-^, *Casp1/11/12*^-/-^), we included mice deficient in type I interferon signaling (*Ifnar1*-/-), which has been shown to be induced via cytosolic nucleic acid sensors RIG-I or MDA5 during blood stage^44^ and hepatic^45^ *Plasmodium* infection. We also included a group of mice lacking γδ T cells (*Tcrd*^-/-^), which have been previously shown to support CD8^+^ T cell activation after SPZ immunization, partly through IL-4 production^13,46^. Consistent with our *in vitro* findings, PbT-I cell expansion was significantly reduced in *Myd88*^-/-^ mice, but not in mice lacking inflammasome components (Fig. 6d). Since MyD88 mediates both TLR and IL-1/IL-18 receptor signaling^47^, the observed defect could involve either pathway. However, because *Il1r1^-/-^* and *Il18r1*^-/-^ mice mounted intact CD8^+^ T cell responses, the defect is more likely attributable to impaired TLR signaling. We also confirmed the requirement for γδ T cells in promoting full CD8^+^ T cell activation in response to sporozoite immunization (Fig. 6d).

To further dissect the relationship between CT, MyD88 signaling, and γδ T cell involvement, we immunized *Myd88*⁻^/^⁻ mice or wild-type mice treated with the γδ-TCR–blocking antibody GL3 with either wild-type or traversal-deficient *Pb*SPZ and measured CD8^+^ T cell responses. In *Myd88*⁻^/^⁻ mice, both the absence of MyD88 (F(1,33) = 45.4, p < 0.001) and the lack of sporozoite traversal (F(1,33) = 55.5, p < 0.001) independently reduced CD8^+^ T cell responses, indicating that the defect in adaptive immunity caused by traversal deficiency is independent of MyD88 signaling (Fig. 6e,f). In contrast, in GL3-treated mice, both γδ-TCR blockade (F(1,33) = 116, p < 0.001) and CT-deficiency (F(1,33) = 11.9, p = 0.0015) also significantly impaired CD8^+^ T cell responses. However, in the presence of GL3, traversal-deficient *Pb*SPZ elicited responses comparable to wild-type parasites, suggesting that the impact of traversal on CD8^+^ T cell priming is mediated through γδ T cells (Fig. 6g,h). Nonetheless, the response to *spect1Δ Pb*SPZ was significantly lower in GL3-treated mice compared to wild-type (p < 0.001), indicating that γδ T cells likely sense additional signals beyond CT.

Finally, we asked whether early γδ T cell activation is impaired in wild-type mice immunized with traversal-deficient *Pb*SPZ. We measured early activation markers on splenic γδ T cell subsets 24 hours post-immunization (Fig. 6i). Activation of Vγ4^+^ and – though to a lesser degree - Vγ1^+^, γδ T cells was reduced in mice immunized with *spect1Δ* parasites compared to controls (Fig. 6j). The partial reduction suggests that γδ T cells respond not only to CT but also to additional sporozoite-derived signals. Taken together, these findings suggest that CT promotes γδ T cell activation, which in turn amplifies CD8^+^ T cell priming following sporozoite immunization.

## Discussion

Vaccination with live SPZ has long been known to offer sterilizing immunity and mechanisms and correlates of protection have been extensively studied^2,6^. Here, we established a reductionist approach that utilizes flow sorting to purify parasites, and profiled innate immune responses induced by SPZ in comparison to those induced by blood stage parasites and LPS. Remarkably, we found that blood stage and SPZ induced different responses and that the response to SPZ overlapped significantly between the two species. Further analysis of expression patterns revealed a unique response to plasma membrane damage induced by SPZ. Consequently, we found that innate cells co-cultured with SPZ unable to induce traversal associated membrane damage showed reduced activation *in vitro*. Importantly, mice immunized with traversal deficient parasite lines showed a defect in CD8^+^ T-cell priming and activation, which likely occurs downstream of γδ T-cell activation. We thus discovered CT-associated membrane damage as a necessary signal for efficient induction of protective cellular immunity by sporozoite immunization.

A long-standing observation in the field has been the explicit necessity of SPZ to be viable during vaccination to achieve protective immunity, a fact that has had severe implications for scaling of the SPZ vaccine. Indeed, viability has been implicated for the efficacy of many other vaccines, such as MMR^48^, with many of the most successful vaccines being live attenuated forms of the original pathogens which often provide live-long protection^49^. Mechanisms such as sensing of mRNA^50,51^ that is only present in viable pathogens have been implicated as checkpoints for the innate immune system to gauge the threat level imposed by a pathogen^52,53^. From our study, membrane damage caused by SPZ traversing through cells, represents an additional viability-dependent signal that is detected by the innate immune system. Our data also contextualize earlier findings of vaccination with chemically attenuated *Pb*SPZ (early liver stage arresting) offering protection^54^ while heat-killed *Pb*SPZ did not offer protection^55^, a phenotype that had been traced to their decreased ability to induce effective CD8^+^ T cell responses^55^.

Taken together, our study illuminates that SPZ traversal-caused membrane damage is a key signal that is picked up by innate immune cells and is relayed through γδ T cells to induce protective CD8^+^ T cell responses. However, while we identified host-CT as a requirement for effective induction of CD8^+^T cell responses, we did not identify a clear sensory pathway which is a limitation of this study. Future work will shed light on the mechanism that senses traversal-induced membrane damage and how this signal is relayed through γδ T cells to help pave the way towards urgently needed, more efficient Malaria vaccines.

## Acknowledgements

We thank support from H. Vohra and M. Devoy of the imaging and cytometry facility at the Australian National University. We acknowledge the instruments and expertise of the Biomedical Resource Facility, the Australian National University, a facility enabled by NCRIS and university support. We thank Benedikt Obermayer-Wasserscheid of the Core Unit Bioinformatics (CUBI) at Berlin Institute of Health (BIH) for single cell RNA-seq raw data pre-processing and Lisa Buchauer for insightful discussions regarding scRNA-seq data analysis. We also thank Calvin Hon for video editing.

## Author contribution

Conceptualization: I.A.C., L.E.S., K.G.P.

Formal analysis: K.G.P., I.A.C., P.M., X.G.

Funding acquisition: I.A.C., L.E.S.

Supervision: I.A.C., L.E.S., F.K., L.B., S.M.M.

Investigation: K.G.P., X.G., J.M., P.M., A.L., L.L., S.L., P.C., C.N., S.B.

Methodology: K.G.P., I.A.C., J.M., H.J.S., A.H., S.B., F.K., L.E.S., L.B.

Visualization: K.G.P., I.A.C., P.M., X.G.

Resources: R.A., L.H., L.Lo., L.B. S.M.M.

Writing – original draft: K.G.P., I.A.C.

Writing – review and editing: K.G.P., I.A.C.

## Methods

### Mouse strains

C57BL/6Crl were used as wild type mice for all experiments. Mice were bred at the Australian Phenomics Facility or at the Department of Microbiology and Immunology and the University of Melbourne. Transgenic strains for innate sensing pathways were: *MyD88*^-/-^(MGI: 2385681) ^56^, *Nlrp3*^-/-^ (MGI: 5465108)^57^, *Aim2*^-/-^ (MGI: 5428935)^58^, *Caspase1/11*^-/-^ (MGI: J:24258)^59^, and *Caspase1/11/12*^-/-^ (gift from Marco Herold, WEHI, Melbourne)^60^. Downstream effector and receptor deficient mice strains were: *Il18*^-/-^ (Obtained from The Jackson laboratory, strain ID #:004130), *IL-1R^-/-^* (gift from Ian Wicks, WEHI, Melbourne) and *Ifnar2*^-/-^ (gift from Paul Herzog, Monash University)^61^. For experiments with antigen-specific CD8 T cells, we used the previously described transgenic PB-T1 strain that was generated using V(D)J segments of the TCRα- and β-genes of a CD8+ T cell hybridoma line reactive to *P. berghei* lysate. The T-cells were later shown to recognize NVFDFNNL derived from the putative *P. berghei* 60S ribosomal protein L6^62^.

### Parasites

All rodent *Plasmodium* parasites were generated in a *P. berghei* ANKA parent line. The *P. berghei* ConF^14^ (RMgm-136) strain which has GFP under the control of the HSP70 promoter was used as the control for all experiments. The *spect1Δ*^14^ (RMgm-139) is the result of a crossing between the ConF parasite and knockout parasites lacking SPECT1^63^ (RMgm-138). The *plp1Δ*^14^ (RMgm-137) is the result of a crossing between the ConF parasite and knockout parasites lacking PLP1^64^ (RMgm-135). The *CelTOSΔ*^14^ is the result of a crossing between the ConF parasite and a “conditional” KO has been generated^65^ in which the 5’UTR of *CelTOS* is replaced by the promoter region of *WARP* (von Willebrand factor A domain-related protein). This parasite is used because CelTOS is also required for transmission stages of the Plasmodium parasites. In this parasite, the use of the WARP promoter restricts *CelTOS* gene expression to the ookinete stage and so near normal numbers of CelTOS-deficient salivary gland sporozoites exhibiting impaired CT activity can be obtained^65^.

NF54-CGL *P. falciparum* sporozoites were obtained from TropIQ Health Sciences, Nijmwegen, NL. The line was created as described before^23^ expressing a GFP-Luciferace fusion protein under the control of the CSP promoter at the *pf47* locus. The 3D7 GFP^+^ parasite line used for blood stage experiments expressed GFP under the control of the Ef1α5 promoter integrated at the *Mal1_18s* locus.

### Primary human cells

Primary human cells were obtained from buffy coats from the German Red Cross Blood Transfusion Service, Berlin, Germany. Information about age and sex of buffy coat donors is not available in accordance with German privacy laws.

## METHOD DETAILS

### Parasite enrichment for co-culture

*P. berghei* sporozoites were dissected from mosquito salivary glands using a binocular fitted with a GFP filter set to GFP^+^ SPZ for a clean dissection. Salivary glands were collected in cold sterile PBS and gently mashed with a pestle inside of a low-adherence tube and passed through a 70 μM cell strainer (BD Biosciences). For *P. berghei* blood stage parasites, blood from infected mice was drawn and diluted 1/1000 in PBS with 2 mM EDTA. Parasites were sorted on a BD Aria II or BD Fusion system using a 100 μM nozzle based on transgenic GFP expression.

*P. falciparum* sporozoites were dissected from *Anopheles stephensi* salivary glands and kept on ice for overnight shipment. The next day, PfSPZ were sorted on a BD FACSMelody based on transgenic GFP expression. Blood stage *P. falciparum* parasites were cultured as previously described^66^. Trophozoite-stage infected erythrocytes were enriched from *in vitro* cultures using magnetic separation on MACS LS columns (Miltenyi Biotec) as described before^67^. In short, MACS LS columns (Miltenyi) were placed into a 4-fold magnetic stand (Miltenyi) and rinsed with sterile PBS. Then, around 15 ml of blood stage culture at 1 % hematocrit and 1 to 30 % parasitemia was added sequentially to each column. Columns were then washed twice with 5 ml sterile PBS and removed from the magnetic stand. Parasites were eluted from the columns with 5 ml sterile PBS containing 2 mM EDTA. Next, parasites were washed and resuspended in PBS. Concentration of purified parasites was assessed using Neubauer hemocytometers while parasitemia was measured with Giemsa smears. Parasitemia of purified blood stage culture was usually between 95 and 99 % with almost exclusively late-stage parasites present. MOI for co-culture experiments was calculated based on batch purity and cell counts. Uninfected erythrocytes from the same RBC donor served as controls.

### BMDM co-culture experiments

Primary bone marrow–derived macrophages (BMDMs) were generated by culturing bone marrow cells from C57BL/6 mice in DMEM supplemented with 10% FCS, 1% Penicillin/Streptomycin, and 30% L929-conditioned medium for 5–6 days. BMDMs were seeded in 96-well plates at 1×10⁵ cells per well in fresh medium without L929 supernatant. GFP^+^ *P. berghei* sporozoites or blood stage parasites were sorted directly onto BMDMs at indicated MOIs. Next, plates were spun down at 300 rcf for 10 minutes and medium was carefully replaced with 100 μl DMEM 10% FCS, 1% P/S. Where indicated, LPS (500 ng/ml) was used as a positive control. After incubation as indicated at 37 °C and 5 % CO2, supernatants were collected for cytokine measurement, and cells were processed for flow cytometry, imaging cytometry, or FACS sorting. For re-cultured cells from RNAseq experiments, GFP^+^ or GFP^-^ cells from all conditions were directly sorted into wells of a 96 well plate containing DMEM with 10 % FCS and 1 % P/S, spun down at 300 rcf for 10 minutes and incubated for 20 h.

### PBMC co-culture experiments

Human PBMCs were isolated by density gradient centrifugation and depleted of T and B cells using magnetic negative selection (Miltenyi Biotec). In short, cells were incubated with CD19 and CD3 MicroBeads (Miltenyi Biotec) according to manufacturers’ instructions for 20 minutes at 4 °C and then passed through a LS column (Miltenyi Biotech) placed in a magnetic stand. Flow-through cells were collected, washed, counted and resuspended in RPMI 1640 (Gibco, ThermoFisher) supplemented with 10% heat-inactivated (HI)FCS, 1% P/S, 1% HEPES, 1% NEAA, 1% GlutaMAX (all Gibco, Thermo Fisher). Next, cells were seeded at 2×10^5^ cells per well in 96-well flat-bottom tissue culture treated plates (Corning) and rested over night before stimulation. Sorted GFP^+^ *P. falciparum* SPZ and enriched blood stage parasites were added at MOIs of 0.5 and 5, respectively. Co-cultures were incubated for 4 hours at 37°C, 5% CO₂. Next, cells were harvested in PBS with 2 mM EDTA and 2 % FCS, stained with anti-human HLA-DR BV421 (Biolegend), CD19 & CD3 PerCP and condition-specific TotalSeq-A antibodies (Biolegend) for 30 minutes at 4 °C. Next, cells were washed 3 times, stained with 7AAD and sorted on a FACSMelody (BD Biosciences).

### Cytokine measurements

Cytokine concentrations from the BMDM cell culture supernatants were quantified using a 4-plex ELISA for TNF, IL-6, CXCL1, and IL-1β (MCYTOMAG-70K, Merck) according to the manufacturers’ instructions. CXCL2 was measured using a DuoSet ELISA kit (R&D Systems) according to manufacturers’ instructions. O-Phenylenediamine (Sigma) was used as HRP substrate, and the reactions were stopped with 1% (v/w) sodium dodecyl sulphate. Absorption was read using a VICTOR Nivo plate reader (PerkinElmer) at 450 nm. Standard curves and biological samples were run in technical duplicates which were averaged during analysis.

### Imaging cytometry

Single cell suspensions from BMDM co-culture experiments were spun down for 5 minutes at 300 rcf and resuspended in 100 µl FACS buffer. Before analysis, 1 µl 7AAD was added to the cells. Cells were then analyzed on a Amnis ImageStream MKII according to the manufacturer’s instructions. Data was analyzed with Amnis IDEAS software. Sporozoite internalization by macrophages was identified using the internalization wizard.

### Live cell imaging

BMDMs were co-cultured with *Pb*SPZ for 15 minutes and then imaged on a Zeiss Axio Observer 7 microscope fitted with a humidified culture chamber kept at 37°C and 5 % CO2. Images were analyzed in FIJI (ImageJ 1.54p) and final edits were done in adobe after effects.

### Rhodamine dextran membrane disruption assay

To assess membrane integrity following SPZ exposure, BMDMs were seeded at 1 × 10⁵ cells per well in DMEM supplemented with 10% FCS, 1% P/S in 96-well plates one day prior to the assay. On the day of the experiment, culture medium was replaced and sporozoites were added at a multiplicity of infection (MOI) of 0.3–0.4. After 1 hour of incubation at 37 °C to allow sporozoite activation after being sorted, Rhodamine Dextran (lysine-fixable, 70 kDa; ThermoFisher) was added directly to the wells at a final dilution of 1:1000. Cells were incubated for 15 minutes at 37 °C, 5% CO2 then harvested and washed once in PBS and immediately analyzed by flow cytometry.

### SmartSeq2-based bulk RNA sequencing

SmartSeq2^68^ was adapted to generate bulk transcriptomes of 50 sorted macrophages. To account for bulk processing of 50 sorted macrophages per well, lysis buffer conditions were initially optimized. Optimal lysis buffer consisted of 1% w/v tween20, 5 mM DTT, 2U/µl RNase inhibitor (Clonetech). Rigid 96 well PCR plates (ThermoFisher) containing 2 µl modified SmartSeq2 lysis buffer per well were prepared and kept on ice until use. Macrophages were washed from the wells with FACS buffer and sorted after adding 7AAD. Macrophages were gated as 7AAD negative single cells and GFP^-^ as well as GFP^+^ populations were sorted into different wells. Per condition and population, 3 to 6 technical replicates of 50 cells were sorted into different wells that were later pooled during analysis. Subsequent reverse transcription, amplification and sequencing followed the smartSeq2 protocol with 22 cycles of PCR amplification. cDNA libraries were sequenced on a NextSeq500 (Illumina) using paired end 75 bp reads

### Single cell RNA sequencing

Sorted cells from co-culture experiments were placed on ice, washed with PBS, counted and pooled across donors. Next, single cell libraries were generated using a 10x GEX 3’ v3.1 kits while loading GFP^+^ and GFP^-^ fractions of cells onto different lanes for later identification. HTO and GEX libraries were generated according to manufacturers’ instructions and sequenced on a S4 flow cell on a NovaSeq6000 (Illumina).

### Mouse immunization experiments

Recipient mice were intravenously injected with 3-6 ×10^4^ irradiated (15kRad) SPZ resuspended in PBS. For peripheral blood mononuclear cell (PBMC) analysis, blood was collected via tail bleed into EDTA-coated tubes. Serum was separated and stored for future ELISA analysis, while remaining red blood cells were lysed for flow cytometry. For spleen analysis, spleens were collected and smashed against a 70 μm cell strainer. The resulting splenocytes were subjected to RBC lysis, stained with an antibody cocktail, and analyzed by flow cytometry. For liver analysis, livers were harvested and mechanically dissociated using a metal mesh, followed by filtration through a 70 μm cell strainer. The resulting suspension was layered over 35% Percoll and subjected to gradient centrifugation. The supernatant was discarded, and the remaining cells were RBC lysed and processed for flow cytometry.

### PbT1 cells

Spleens were harvested from PbT-I transgenic mice (CD45.2^+^) and passed through a 70 µm strainer to obtain single-cell suspensions. Red blood cells were lysed and splenocytes were counted using a hemocytometer. Cells were labelled with CellTrace Violet (CTV; Invitrogen) and 2×10⁶ cells were injected intravenously into congenic CD45.1^+^ recipient mice.

### Antibody ELISA

Antibodies targeting P. berghei CSP in immunized or naïve mice were measured using in-house developed ELISAs. In short, Nunc MaxiSorp plates (ThermoFisher) were coated overnight with 1 µg/ml streptavidin (Sigma) in PBS. On the next day, plates were washed with ELISA washing buffer, blocked for 1 h in ELISA blocking buffer and incubated with biotinylated recombinant *P. berghei* CSP (1 µg/ml in PBS, Genscript) for 1 h at room temperature. Next, plates were thoroughly washed and incubated with sera collected from immunized mice in serial dilutions for 2 h at room temperature. After another washing step, HRP anti-mouse total IgG or HRP anti-mouse IgM (both Southern Biotech) antibodies were added to the wells (1 in 10000 dilution) for 1 h. Next, plates were washed 5 times and developed using KPL peroxidase components A and B (SeraCare). Reaction was stopped with 1% (v/w) sodium dodecyl sulphate and read at 450 nm on a VICTOR Nivo plate reader. Antibody concentrations were calculated as area under the dilution curve (AUC).

### Bulk RNA-seq data analysis

Quality control of raw sequencing data was performed using FastQC. For mouse bulk RNA sequencing data, reads were aligned to mouse reference genome GRCm38.98 using Kallisto^69^ with 30 bootstraps. Technical replicates were merged at this step. Differential gene expression was analyzed using DEseq^70^ with gene count output of Kallisto imported using tximeta^71^. Volcano plots of differentially expressed genes were generated using EnhancedVolcano^72^ and functional analysis was carried out using ClusterProfiler^73^ and fgsea.

### scRNA-seq data analysis

Raw sequencing reads were aligned to a custom reference genome built with *Homo sapiens* (GRCh38) and *Plasmodium falciparum* FASTA and GTF using Cell Ranger v5.0.0 (10x Genomics). The pipeline further handled sample demultiplexing, identification of cell barcodes and gene expression quantification. Next, to remove technical artifacts such as ambient RNA, CellBender^74^ was applied to the raw feature matrices produced by Cell Ranger. Donor demultiplexing was carried out using vireoSNP^75^ (v0.5.9), using information on allele counts per cell for known and common SNPs obtained from cellSNP^76^ (v0.3.0). Seurat v5^77^ was used to process each library. Cells were filtered based on number of UMIs per cell, number of detected genes per cell and the percentage of mitochondrial gene expression per cell. Each library was then normalized, scaled and subjected to dimensionality reduction. Next, they were integrated using IntegrateLayers() with “CCAIntegration” method. Cell clusters were then defined with FindNeighbors() and FindClusters(). Where applicable, resolution (parameter “res”) was kept uniform at 1 and PCA dimensions 1 through 30 were chosen (dims = 1:30). Cell clusters were manually annotated based on canonical markers, and contaminating T cells, B cells, and dendritic cells were removed.

For differential expression (DE) analysis, pseudobulk counts were generated using AggregateExpression() and DE genes were detected with FindMarkers() and test.use = “DESeq2”. Gene set enrichment analysis was performed using fgsea package^78^ (v1.34.0) and gseGO() function in clusterProfiler package (v4.16.0). Module scores or plasma membrane injury gene set were computed using Seurat’s AddModuleScore().

### Statistical analysis and visualization

Further visualization and data exploration were supported by EnhancedVolcano (v1.26.0), UpSetR (v1.4.0) and ComplexHeatmap (v2.22.0) packages in R. GraphPad Prism (v.10.4.2) was also used for plotting and statistical analysis and figures were generated in Adobe Illustrator (29.3.1). (v4.16.0). Module scores or plasma membrane injury gene set were computed using Seurat’s AddModuleScore().

### Ethics statement

All animal procedures were approved by the Animal Experimentation Ethics Committee of the Australian National University (Protocol number: 2019/36 and 2022/36). All research involving animals was conducted in accordance with the National Health and Medical Research Council’s Australian Code for the Care and Use of Animals for Scientific Purposes and the Australian Capital Territory Animal Welfare Act 1992. Permission for experiments with human primary cells from buffy coats was obtained from Charité Ethics Committee (Protocol number: EA1/004/12, Charité – University Medical Centre, Berlin, Germany).

### Data availability

Bulk RNA-seq data has been deposited to the European Nucleotide Archive (ENA) and is publicly available with accession number PRJEB90938. Single cell RNA-seq count matrices are available on Gene Expression Omnibus (GEO) under accession number GSE301430. Raw sequencing data could not be uploaded in accordance with German and European privacy regulations.

### Code availability

No custom algorithms were used for data analysis. The code generated to analyze RNA-seq data is made available on GitHub at github.com/KaiP1/Pohl-et-al.-DEseq2-analysis-pipeline.git.

