## Supplementary Figures and Tables for "Innate immune sensing of cell traversal by *Plasmodium* sporozoites drives protective T cell responses"

### Extended data

#### Extended data Fig.1: Co-culture of P. berghei infected RBCs with BMDMs.

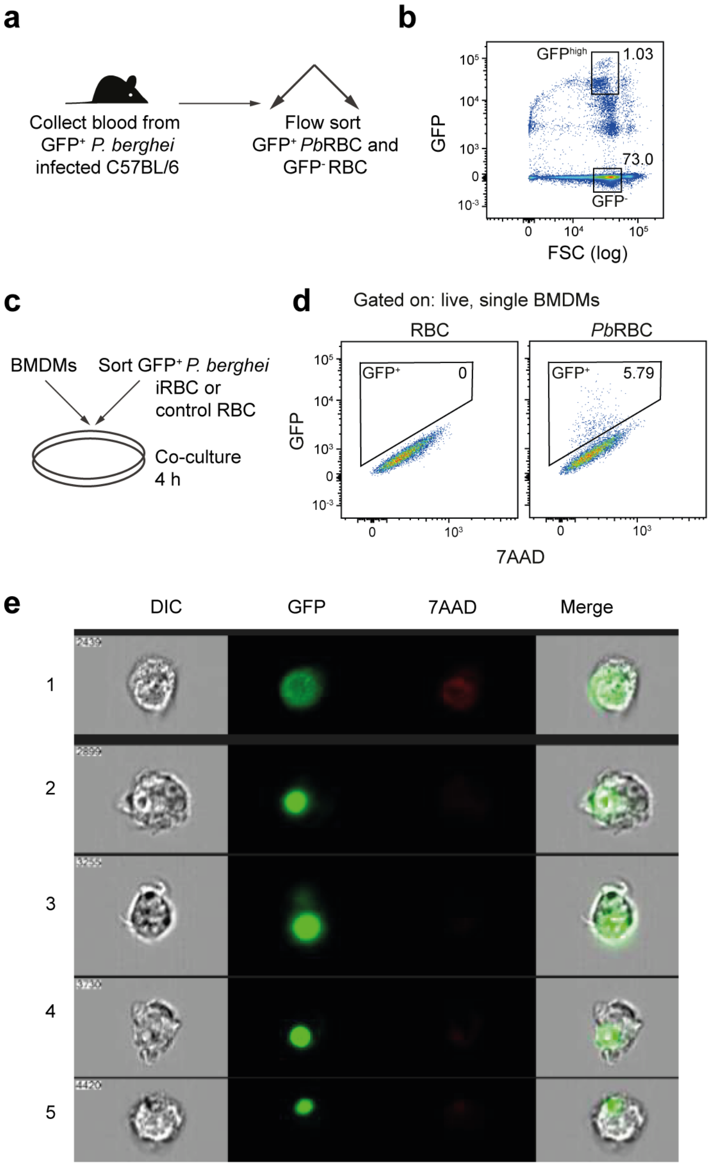


**a,** Schematic of experimental setup. **b,** GFP$^+$ *Pb*RBCs are clearly detectable in blood from infected mice. **c,** Schematic of experimental setup. **d,** A population of GFP$^+$ macrophages is present after 4 h of co-culture with *Pb*RBC by flow cytometry. **e,** Imaging cytometry of GFP$^+$ macrophages after 4 h of co-culture with *Pb*RBC.

### Extended data Fig. 2: Transcriptional profiling of *Plasmodium*-exposed BMDMs

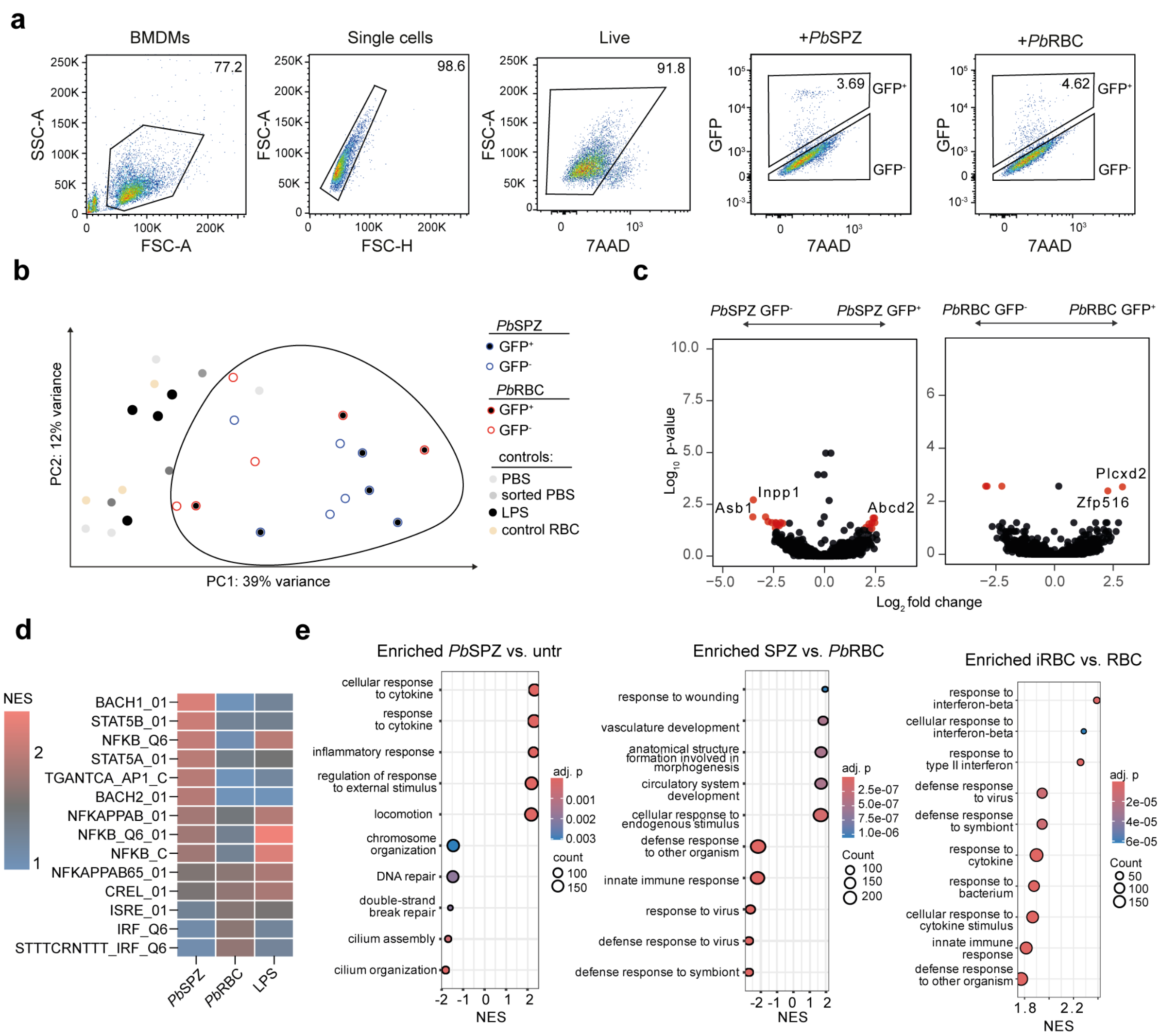


**a,** Gating strategy to sort GFP+ and GFP- BMDMs. **b,** PCA of BMDMs after 4 h of stimulation as indicated followed by sorting of GFP+ and GFP- BMDMs and re-culture for 20 h shows similar transcriptomic signatures upon treatment with either life cycle stage. **c,** Volcano plots showing DEGs of GFP+ vs. GFP- BMDMs after co-culture with *Pb*SPZ or *Pb*RBC for 4 h. **d,** GSEA for transcription factor target sites around regulated genes. NES for top enriched transcription factors shown ordered by NES for BMDMs stimulated with *Pb*SPZ. **e,** GSEA for GO BPs on genes ranked by $log_2$ fold change in the indicated comparisons.

**Extendend data Fig. 3:In vitro investigation of human primary immune cell interactions with *P. falciparum* SPZ and blood stage**

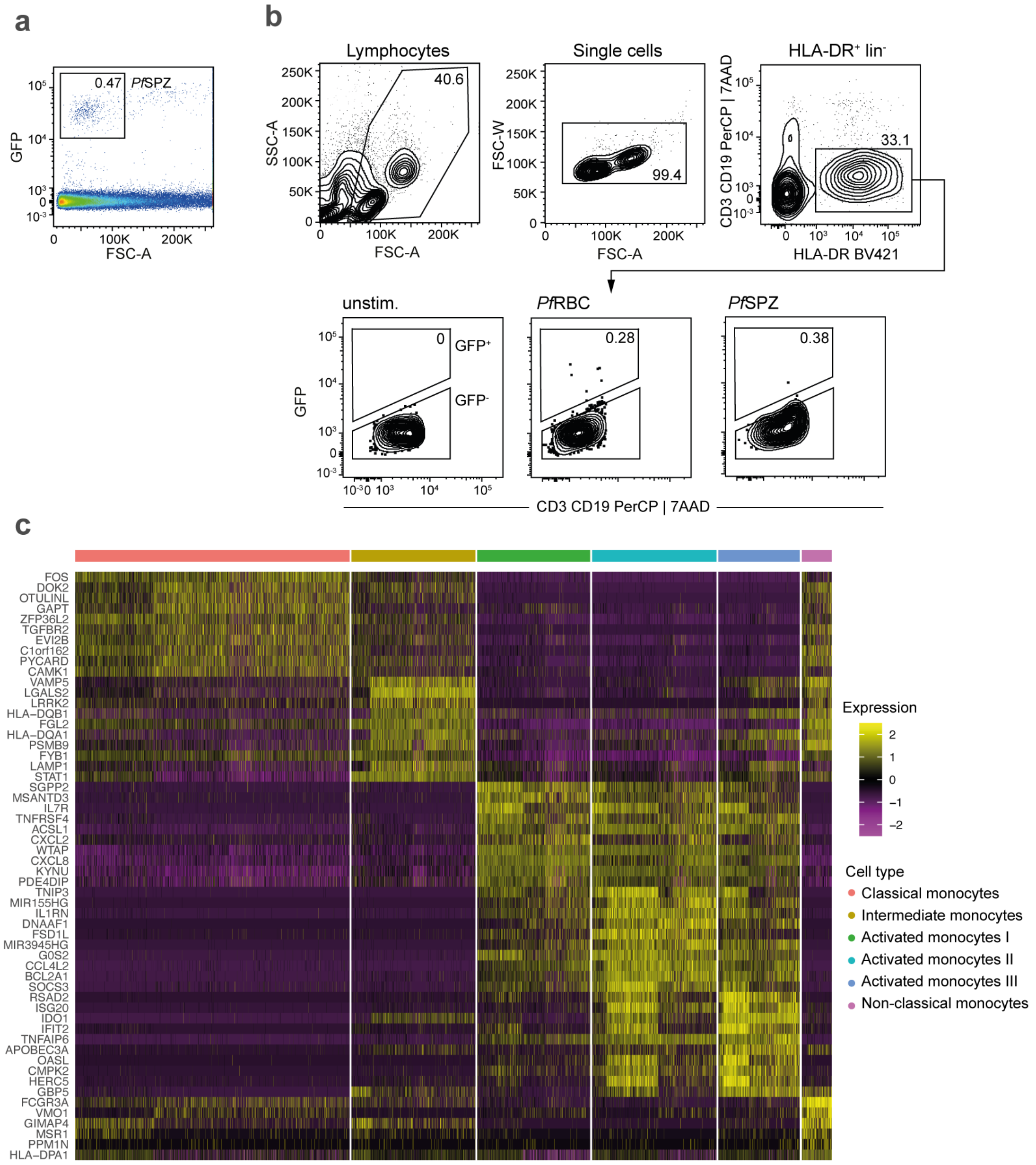


**a,** GFP$^{+}$ NF54-CGL *P. falciparum* SPZ are readily detectable in salivary gland extracts. **b,** Gating strategy used to sort GFP$^{+}$ and GFP$^{-}$ primary human innate immune cells after co-culture with *Pf*RBC, *Pf*SPZ, or control RBC. **c,** Heatmap depicting marker genes for the identified cell types.

### Extended data Fig. 4: Innate activation induced by PfSPZ and PfRBC.

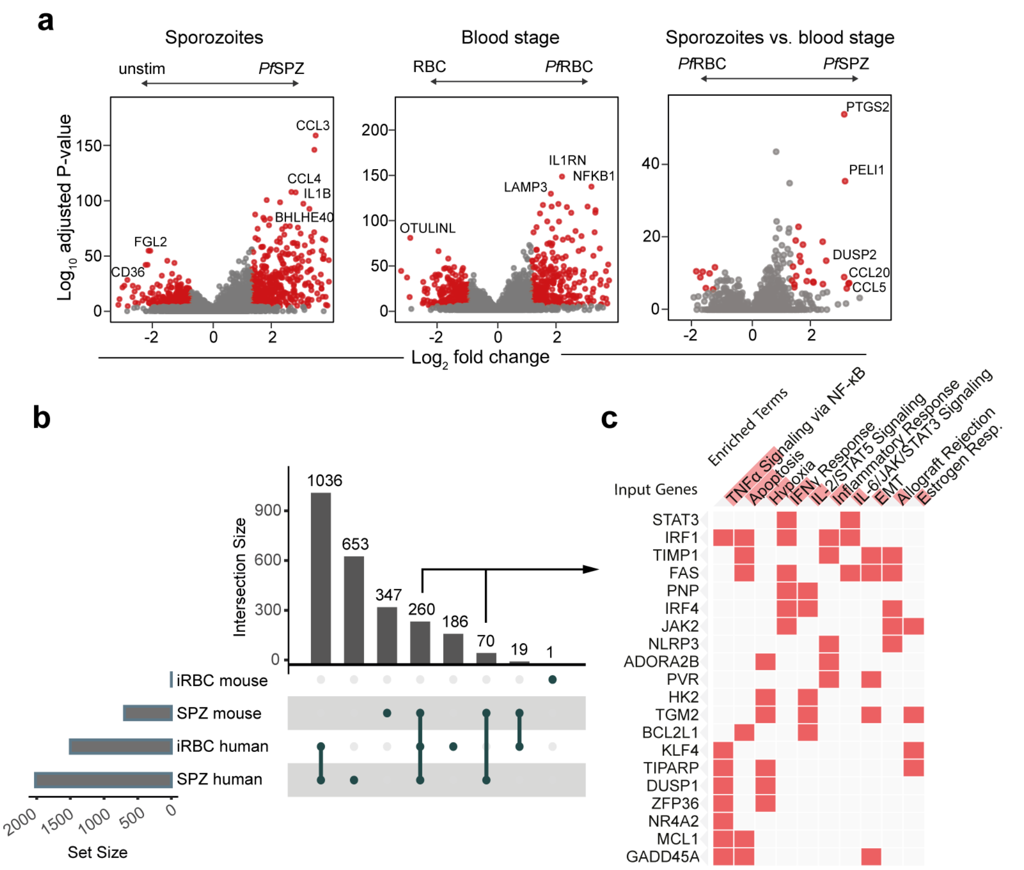


**a,** Volcano plots showing DEGs between indicated treatment groups, calculated using DESeq2 after pseudobulking of scRNAseq data. **b,** Upset plot of upregulated genes ($log_2$ fold change > 1.5, padjust < 0.05) across conditions and host-parasite pairs showing substantial overlap between SPZ stimulated mouse and human innate cells. **c,** Overrepresentation analysis of genes overlapping between SPZ stimulated human and mouse innate cells.

**Extended data Fig. 5: Activation and wounding of BMDMs by PbSPZ.**

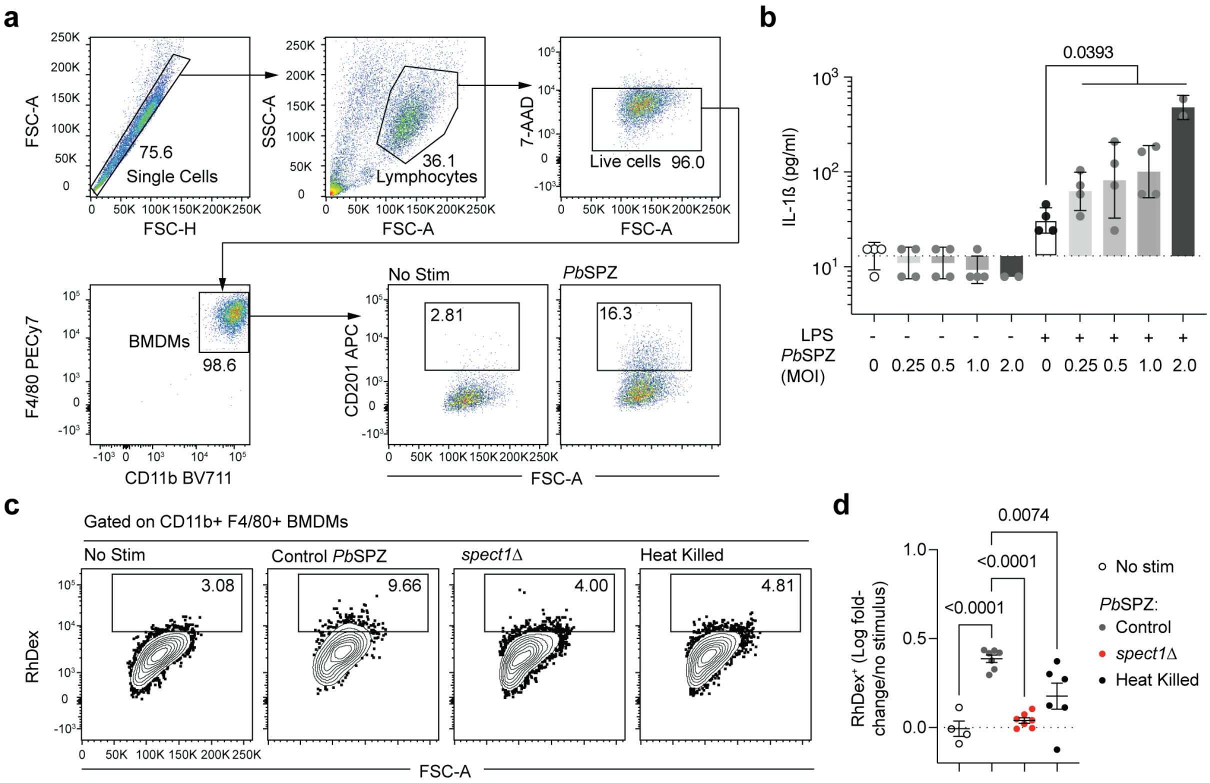


**a,** Gating strategy for mature BMDMs in culture and detection of CD201 on the cell surface. **b,** IL-1β levels in the supernatant after 24 hours of co-culture with *Pb*SPZ at the indicated MoIs, w/wo priming with LPS; data shown are biological replicates from 2 independent experiments, mean ± SD shown, analysis via linear mixed model. **c,** Representative flow cytometry plots showing uptake of RhDex after 15 minutes of co-culture of BMDMs with *Pb*SPZ. **d,** Quantification of data from **c**; data are replicates from a single experiment, mean ± SD shown, analysis via one-way ANOVA with Tukey's post-test.

**Extended data Fig. 6: Analysis of adaptive immune responses to CT deficient PbSPZ**

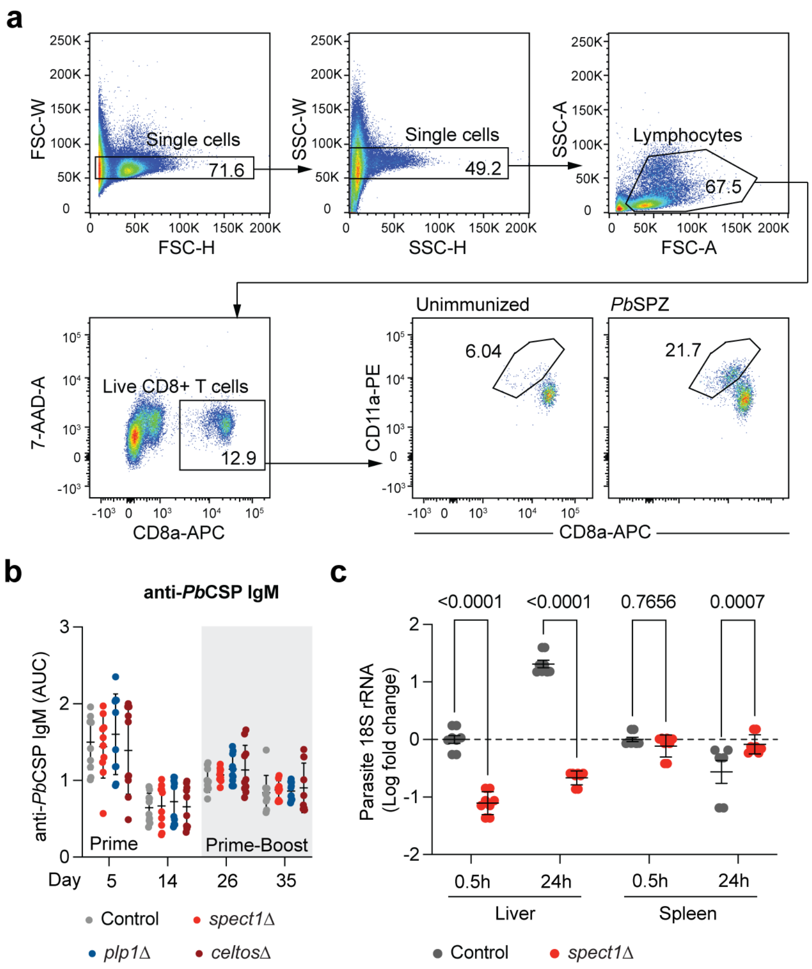


**a,** Gating strategy for identification of CD8$^+$ T cells in the blood of immunized mice; similar gating was used for the identification of CD8$^+$ T cells from the spleen and liver. **b,** Anti-*Pb*CSP IgM in the sera of mice immunized as in Fig. 5 A; data pooled from 2 independent experiments, mean ± SD shown, analysis via linear mixed model, no significant differences were observed across any groups. **c,** RT-PCR analysis of *P. berghei* 18s rrRNA in the liver and spleen at the indicated times post infections; data from 2 independent experiments and are presented as fold change relative to the liver at 30 minutes; mean ± SD shown, analysis via linear mixed model.

### Supplementary table 1: Antibodies

| Antibody | Source | Identifier |
|---|---|---|
| Anti-mouse CD201 APC | Biolegend | #Cat: 141505, Clone: RCR-16 |
| Anti-mouse CD8a APC | Biolegend | #Cat: 100712 Clone: 53-6.7 |
| Anti-mouse CD11a PE | Biolegend | #Cat: 101107 Clone: M17/4 |
| Anti-mouse CD8a APC | Biolegend | #Cat: 100712 Clone: 53-6.7 |
| Anti-mouse CD11a PE | Biolegend | #Cat: 101107 Clone: M17/4 |
| Anti-mouse/human CD44 APC | Biolegend | #Cat: 103028 Clone: IM7 |
| Anti-mouse CD62L BV711 | Biolegend | #Cat: 104445 Clone: MEL-14 |
| Anti-mouse CD69 APC | Biolegend | #Cat: 104514 Clone: H1.2F3 |
| Anti-mouse CD25 APC/Cyanine7 | Biolegend | #Cat: 102026 Clone: PC61 |
| Anti-mouse γδ T-Cell Receptor BUV395 | BD | #Cat: 744118 Clone: GL3 |
| Anti-mouse CD3 PerCP/Cyanine5.5 | Biolegend | #Cat: 100218 Clone: 17A2 |
| Anti-mouse TCR Vγ1.1 BV605 | BD | #Cat: 745144 Clone: 2.11 |
| Anti-mouse TCR Vγ4 PE | BD | #Cat: 569158 Clone: 49.2 |
| Anti-mouse TCR Vα8.3 FITC | Biolegend | #Cat: 127705 Clone: B21.14 |
| Anti-mouse CD45.1 PE | Biolegend | #Cat: 110708 Clone: A20 |
| Anti-human HLA-DR BV421 | Biolegend | #Cat: 307635 Clone: L243 |
| Anti-human CD19 PerCP | Biolegend | #Cat: 302227 Clone: HIB19 |
| Anti-human CD3 PerCP | Biolegend | #Cat: 300427 Clone: UCHT1 |

### Supplementary table 2: Mouse and parasite strains

| Mouse/Parasite strain | Source | Identifier |
| --- | --- | --- |
| *P. berghei* ConF GFP (parasite) | Amino et al.[14] | RMgm-136 |
| *P. berghei* ConF GFP spect1$^{-/-}$ (parasite) | Amino et al.[14] | RMgm-139 |
| *P. berghei* ConF GFP *CelTos*$^{-/-}$ (parasite) | Amino et al.[14] | - |
| *P. berghei* ConF GFP plp1$^{-/-}$ (parasite) | Amino et al.[14] | RMgm-137 |
| *P. falciparum* NF54 GFP-luc (parasite) | Miglianico et al.[23] | - |
| *P. falciparum* 3D7 GFP (parasite) | Alexander Maier (ANU, RSB) | - |
| C57BL/6Crl (mouse) | The Jackson laboratory | MGI:2160152 |
| MyD88$^{-/-}$ (mouse) | Adachi et al.[56] | MGI: 2385681 |
| NLRP3$^{-/-}$ (mouse) | Kovarova et al.[57] | MGI: 5465108 |
| Caspase1/11$^{-/-}$ (mouse) | Kuida et al.[59] | MGI: J:24258 |
| Aim2$^{-/-}$ (mouse) | Jones et al.[58] | MGI: 5428935 |
| IL-1R$^{-/-}$ (mouse) | Gift from Ian Wicks (WEHI, Melbourne) | N/A |
| IL18$^{-/-}$ (mouse) | The Jackson laboratory | Strain #:004130 |
| Caspase1/11/12$^{-/-}$ (mouse) | Salvamossa et al.[60] gift from Marco Herold (WEHI, Melbourne) | N/A |
| IFNAR2$^{-/-}$ (mouse) | Hardy et al.[61] | MGI: 2680693 |
| PBT1 (mouse) | Lau et al.[79] | N/A |
